# Developmental temperature drives distinct transcriptomic responses to acute temperatures and correlated differences in thermal tolerance

**DOI:** 10.1101/2025.09.18.676804

**Authors:** Alison Hall, Manali Rege-Colt, Melissa Pespeni

## Abstract

Marine invertebrate populations exhibit varying capacities to withstand rising environmental temperatures, but the genetic basis of this differential tolerance remains an important area of investigation. Plasticity can be a valuable tool in the arsenal of an animal trying to maintain physiological function under rapidly changing conditions and can be an important contributor to thermal tolerance. In this work, we characterized the transcriptomic response to elevated developmental temperature, and subsequent acute exposure to two higher temperatures using the widespread and ecologically important copepod, *Acartia tonsa*. Using a split brood experimental design, we found that copepods that developed at 22°C had higher upper lethal temperatures compared to those reared at 18°C, demonstrating developmental plasticity in thermal tolerance. Transcriptomic analyses revealed that developmental temperature strongly influenced gene expression both at baseline and in response to acute thermal stress. Exposure to a moderate heat challenge (28°C) elicited divergent transcriptional responses between developmental treatments, suggesting developmental preconditioning, whereas exposure to extreme heat (33°C) triggered a more conserved stress response across groups. Weighted gene co-expression network analysis (WGCNA) identified gene modules associated with upper lethal temperature, highlighting stress response, cellular regulation, and metabolic pathways as key contributors to thermal tolerance. Together, our results reveal how developmental environments shape gene expression patterns and thermal phenotypes, providing insight into the molecular basis of plasticity and potential resilience to climate change.

## Introduction

The ocean absorbs 93% of the heat produced by global climate change, and most of that excess heat is stored near the surface (G. C. Johnson & Lyman, 2020). With this increase in mean heat absorption, the frequency of extreme heat anomalies is also rising. The global ocean heat content, measured from the surface to a depth of 2,000 meters, has continued to increase, reaching new record highs in 2025. In many regions, sea surface temperatures have been observed to be up to 2°C above the 20^th^ century average, contributing to widespread marine heatwaves and elevated risks for marine ecosystems (Ncei., 2024). Marine populations must cope with these changes, and their ability to survive in novel conditions is critical to future ocean ecosystem health (Wernberg et al., 2024). Marine invertebrates exhibit varying capacities to withstand rising environmental temperature and understanding the genetic bases of differential. thermal tolerance is crucial for predicting species’ resilience to climate change (Angilletta & Angilletta, 2009; Sunday et al., 2011; K. M. Johnson & Hofmann, 2017; Manullang et al., 2025). Among marine invertebrates, copepods are of particular interest because they are abundant, ecologically critical zooplankton with short generation times, making them ideal models for studying rapid responses to environmental change (Turner, 2004).

Rapid acclimation to novel thermal conditions can be achieved through phenotypic plasticity, and this phenotypic variation can be an important contributor to thermal tolerance (Angilletta & Angilletta, 2009; Lockwood et al., 2010; Logan & Somero, 2011; Fox et al., 2019). Plasticity refers to an organism’s ability to adjust its physiology or behavior in response to environmental changes without a change in DNA composition (Pigliucci, 2005; Scheiner, 1993). Developmental plasticity occurs when environmental conditions experienced during an organism’s development induce changes in its phenotype. Developmental plasticity enhances survival in new environments by allowing environmentally induced changes to align the adult phenotype more closely with its surroundings (Beldade et al., 2011). This type of plasticity can be especially significant in relatively short-lived animals, as it may support acclimatory responses to the variability associated with climate change. Plasticity can play an even more important role in thermal tolerance than genetic variation (Hoffmann et al., 2005; Ashlock et al., 2024). In relatively short-lived species, such as copepods, developmental plasticity may play a critical role in supporting acclimatory responses to rapid environmental changes, such as those associated with global warming. Despite its importance, key gaps remain in our understanding of the genomic mechanisms underlying plasticity. Questions include which genes plastically respond to changes in thermal environments and which specific genes are critical for the upper lethal temperature response.

If a species can mount a plastic physiological response to stress, it may face lower extinction rates and have a greater ability to remain within its native range (Gunderson & Stillman, 2015; Snell-Rood et al., 2018). While plasticity is a key component of many organisms’ responses to environmental stress, the diversity of mechanisms involved and whether they are taxon-specific remain incompletely characterized (Burraco et al., 2017). The gene expression response studied through RNA sequencing, can provide a mechanistic understanding of stress tolerance (Rivera et al., 2021). Recent studies have demonstrated that ocean warming can lead to significant changes in gene expression and increased aerobic demand in marine ectotherms, underscoring the necessity to understand the molecular basis of thermal tolerance (Eaton et al., 2022). Transcriptomics has been identified as a powerful approach to understanding whether plasticity supports an acclimatory response to novel thermal environments in non-model species (DeBiasse & Kelly, 2016).

To fully understand the molecular basis of thermal tolerance, it is important to examine how organisms respond to multiple levels of heat exposure rather than a single threshold. Studies that include a range of acute temperature treatments can reveal whether plastic responses follow a graded pattern or if there is a threshold beyond which a generalized stress response dominates (Logan & Somero, 2011). Investigating transcriptional changes across multiple acute temperatures provides insight into whether plasticity allows fine-tuned acclimation to moderate warming or if extreme heat overrides developmental programming (Gaitán-Espitia et al., 2017; Jeffries et al., 2021). By comparing responses to acute sublethal and more stressful thermal conditions, we can assess whether different temperature exposures elicit different gene expression responses and whether this varies with developmental temperature.

In this study, we investigated differences in gene expression between copepods developed at ambient and elevated temperatures, both at their baseline developmental temperatures and after exposure to two acute, sublethal thermal challenges using RNA sequencing (Fig. 1A). The copepod *Acartia tonsa* was chosen for this study due to its broad distribution, critical role in coastal food webs (Turner, 1984), and previously observed phenotypic plasticity in traits related to thermal tolerance (Healy et al., 2019; M. C. Sasaki & Dam, 2020). We observed that spending a single generation at 22°C results in improved thermal tolerance compared to copepods that developed at 18°C (Fig. 1B), consistent with previous findings of developmental plasticity in this trait (M. Sasaki et al., 2019; Ashlock et al., 2024). To understand the molecular bases of these phenotypic differences, our first objective was to characterize gene expression differences between developmental temperature treatments both at baseline and following acute, sublethal heat exposures (28°C and 33°C), to evaluate how developmental temperature shapes transcriptional responses to thermal stress. Our second objective was to identify functional groups of genes and co-expression modules whose expression correlates with thermal tolerance, to gain insight into the molecular pathways contributing to plasticity and physiological resilience. Third, we aimed to test whether the severity of thermal stress influences the extent to which transcriptomic responses are developmentally programmed or conserved, comparing molecular responses to moderate (28°C) versus extreme (33°C) acute heat. By examining responses at multiple acute temperatures, we aimed to provide a more nuanced understanding of how gene expression patterns shift under different levels of thermal stress. By elucidating differences across treatments and the specific genes and pathways involved, we can better predict how organisms will respond to ongoing environmental changes and inform conservation strategies aimed at preserving marine biodiversity in the face of climate change.

**Figure 1:**
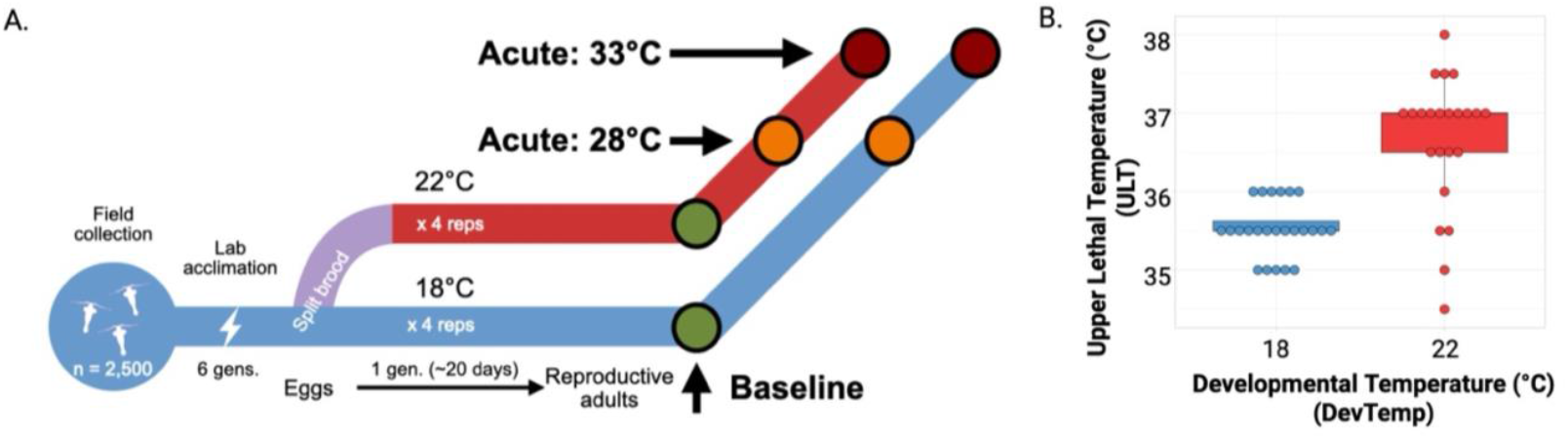
Schematic of the experimental design to assess gene expression responses both within and between two developmental temperature treatments, and across three acute temperature exposures (A). Upper lethal temperature for copepods developed at two developmental temperatures (18°C or 22°C) for a single generation. Each dot represents data for an individual copepod and color represents developmental temperature (18°C in blue or 22°C in red) (B).

## Methods

### Collection and Culture Maintenance

Copepods were collected from Cape May, New Jersey (38.95366, −74.88978) on June 6^th,^ 2022, and transported back to the University of Vermont in temperature-monitored coolers by car. At the University of Vermont, a dissection microscope was used to sort n=2,500 *Acartia tonsa* to initiate experimental cultures. The initial culture was maintained at 30psu (practical salinity units) at 18°C in Thermoscientific incubators on a 12h light/dark cycle for over 6 generations before the experiment began. Cultures were fed *ad libitum* with three species of phytoplankton: a green flagellate (*Tetraselmis* sp.), a brown unicellular diatom (*Thalassiosira weisslogii)*, and a red cryptomonad species (*Rhodonomas salina)*. Phytoplankton were cultured in 30psu salt water with F/2 media at 20°C under 20 hours of light and 4 hours of darkness. A 50% water change was done in each copepod culture weekly.

### Split Brood Design

We employed a split-brood design to test how developmental temperature affects adult thermal tolerance and gene expression responses at baseline conditions, 18°C (ambient) or 22°C (projected +4°C warming; Bindoff et al 2019) and after acute elevated sublethal temperature exposures (28°C and 33°C). Eggs produced by common-gardened adults were split into the two developmental temperature treatment groups 18°C (ambient, hereafter DT18) and 22°C (elevated, hereafter DT22), with 4 replicates of each treatment (Fig. 1A).

### Gene Expression Response to Acute Exposure

Whole, pooled animal RNA-sequencing was used the quantify the molecular responses under three conditions for each of the developmental temperatures: Baseline (BASE: 18°C or 22°C), acute 28°C (A28), and acute 33°C (A33) (Fig. 1A). For Baseline: twenty-five untreated adult females were collected from each replicate and held for 1 hour at their developmental temperature, mirroring the handling time and conditions of acutely stressed animals. For acute sublethal exposures: A separate set of 25 adult females per replicate from both developmental temperatures was divided among five tubes with 6-ml seawater (5 females each) and rested for 1 hour at their developmental temperature. The temperature was then ramped from the developmental temperature (either 18°C or 22°C) to the acute temperature (28°C or 33°C) at 1°C per 5 minutes. Pilot trials confirmed 100% survival after 7 hours at 33°C, validating these challenges as sublethal. After one hour at the target temperature, we confirmed that all animals had survived and collected them from the glass tubes using a 5mL transfer pipette and pooled by replicate (n=25 individuals) into cryovials. Excess seawater was removed; samples were flash-frozen in liquid nitrogen and stored in a −80°C freezer until extraction. In total, 24 samples were saved (2 developmental temperatures x [BASE, 28°C, 33°C] x 4 replicates).

### Upper Lethal Temperature (ULT) phenotype assay

At adulthood, six females from each replicate (n=24 per developmental temperature) were assessed for individual upper lethal temperature. Pairs of females were placed in a 20mL 12.5 × 1.5cm glass tube with 6 ml of artificial seawater and submerged in a programmable water bath (A24B, Thermo Scientific, Waltham, MA, USA). Temperature was increased from the respective developmental temperatures to 30°C at 1°C per 5 minutes. At 30°C and each subsequent 0.5°C increment, the temperature was held for 10 minutes. After 10 minutes, a timer was set for an additional 10 minutes, during which time survival was evaluated. Individuals that failed to respond to light and gentle pipette stimulation of the water around the copepod were scored as dead; the bath temperature at that time was recorded as the ULT (Fig. 1B).

### RNA extraction and sequencing

RNA from the 24 samples was extracted using TRIzol reagent (Invitrogen, Carlsbad, CA, USA) and purified with Qiagen RNeasy spin columns. We quantified RNA using the Qubit RNA Broad Range Assay Kit (ThermoFisher, Waltham, MA, USA). RNASeq libraries were prepared by Novogene (Sacramento, CA, USA) and sequenced with 150bp paired-end reads on Illumina NovoSeq 6000. There were 21 samples with RNA of sufficient quality to be sequenced, and we excluded one sample after observing it as an outlier in PCA leaving us with 20 total samples for analysis of differential gene expression (n = 3 or 4 per treatment group). Full RNA sequencing data are available on the NCBI Sequencing Read Archive (SRA) (Accession number PRJNA1322529).

### Gene expression analysis using DESeq2

Raw transcriptome reads were trimmed for quality using FastP(v. 0.23.0) (Chen et al., 2018). We used FastP to detect and trim adapter sequences and removed reads that were of low quality (Phred score < 20) or too short (length < 35 bases). We used Salmon (v. 2.2.1) to simultaneously map to a previously generated *A. tonsa* reference transcriptome (Jørgensen et al., 2019) and quantify transcript abundance (Patro et al., 2017). We used the R package tximport (v. 3.2.0) to convert transcript abundance to gene level counts and import data into DESeq2 (Soneson et al., 2015). We filtered out genes for which there were counts of less than 10 in more than 75% of total samples, leaving high-quality gene expression data for 35,527 transcripts. Gene names, counts, and log2-fold change for all genes available on Dryad (DOI: 10.5061/dryad.866t1g23z). Counts were normalized and log transformed using DESeq2 (version 1.38.3), (Love et al., 2014). All preprocessing steps prior to differential expression analysis, including read trimming and transcript quantification, were performed on the Vermont Advanced Computing Core (VACC) at the University of Vermont. All subsequent analyses were performed in R version 4.2.3.

We used DESeq2 to identify differentially expressed genes (DEGs) in response to developmental temperature, acute temperature, and their interaction (model: ∼ DevTemp + FinalTemp + DevTemp:FinalTemp). DEGs were identified using DESeq2 by filtering genes with an adjusted *P*-value < 0.05 (Benjamini-Hochberg correction). We transformed the read count data using variance stabilization with the vst() function, then used Principal Component Analysis (PCA) to visualize the relative contributions of developmental temperature and acute temperature exposure to the variation in global gene expression patterns.

To visualize differential gene expression patterns and identify genes of interest for functional enrichment analysis, we generated volcano plots and Euler diagrams for each developmental and acute temperature treatment comparison. Volcano plots were constructed using DESeq2 output to display the log_2_ fold change versus adjusted *P*-value for each gene, highlighting significantly differentially expressed genes (adjusted *P* < 0.05). Venn diagrams were used to identify shared and unique sets of differentially expressed genes between treatment groups, providing insight into the specificity and overlap of transcriptional responses. These gene sets were subsequently used in Gene Ontology (GO) enrichment analyses (described below).

To quantify the similarity in transcriptional responses to acute heat exposure between developmental temperature treatments, we conducted pairwise linear regressions using gene-level log_2_ fold change values. For each gene, we extracted the log_2_ fold change values for DT18 and DT22 separately at each acute exposure. These values were merged by gene ID to create matched datasets of fold changes across developmental treatments. Linear models were then fit using lm() in R, with DT22 fold change as the dependent variable and DT18 fold change as the predictor. The resulting R^2^ values were used to assess the degree of correlation in gene expression responses between developmental treatments at each temperature. We visualized the data as scatterplots using ggplot2, colored genes based on their significance category (differentially expressed in DT18 only, DT22 only, both, or neither). This approach allowed us to visualize and statistically evaluate the extent to which acute transcriptional responses were shared or divergent between developmental treatments under moderate (28°C) and extreme (33°C) stress conditions.

### Functional enrichment analysis

To identify the top biological functions associated with differentially expressed genes (DEGs), we performed Gene Ontology (GO) enrichment analysis using the topGO R package (v. 2.50.0) (Alexa & Rahnenführer, 2009). GO term annotations were obtained from a custom annotation file (Brennan et al., 2025) and filtered to retain genes present in the DEG dataset. GO enrichment analysis was conducted using the “parentChild” algorithm with Fisher’s exact test, which accounts for hierarchical relationships among GO terms, reducing redundancy and highlighting broader, biologically relevant categories. We analyzed the biological process (BP) ontology and filtered enriched terms based on annotation size (15– 500 genes) to exclude categories that had too few members or too many. To account for multiple testing, we applied a false discovery rate (FDR) correction (Benjamini–Hochberg method), and terms with adjusted P < 0.05 were considered significant.

Results were visualized using ggplot2 (v. 3.5.1) (Wickham, 2011) with the top 30 enriched GO terms ranked by significance (-log10 p-value). Point size represented the number of significant genes, and additional metrics included the gene ratio (significant genes/total annotated genes) and the number of annotated genes. Separate visualizations were created for ambient and elevated temperature conditions, enabling a comparative analysis of biological processes associated with thermal stress responses.

### Network analysis using WGCNA

We used weighted gene correlation network analysis (WGCNA) using the R package WGCNA (v. 1.73) to identify highly correlated genes as ‘modules’ and test for an association between correlated gene modules and upper lethal temperature across developmental temperature treatment groups (Langfelder & Horvath, 2008). This analysis focused on samples collected at baseline for each developmental temperature (i.e., no acute temperature exposure) to identify gene modules whose expression variation correlated with phenotype data and upper lethal temperature phenotype data were only relevant to these samples. A signed gene co-expression network was constructed using a soft-thresholding power selected based on the scale-free topology criterion using the pickSoftThreshold() function in WGCNA. We chose 8 which was the lowest power at which the scale-free topology model fit (R^2^) exceeded 0.8, while also preserving sufficient mean connectivity to retain biologically relevant gene-gene relationships.

To visualize gene expression patterns in WGCNA modules associated with upper lethal temperature (ULT), we selected genes from the *green* and *darkgrey* modules, which showed the strongest positive and negative correlations with ULT, respectively. Gene lists for these modules were extracted, and variance-stabilized expression values from DESeq2 were used to calculate mean expression per sample as a summary measure of module activity. These module expression values were compared across developmental and acute temperature treatments using boxplots to evaluate how module-level expression varied with experimental conditions. In addition, module eigengenes were extracted and plotted against ULT to visualize the strength and direction of module–trait associations. Scatterplots with linear regression fits were used to assess correlations between ULT and module eigengenes, providing a summary of how each module’s expression covaried with thermal tolerance.

## Results

### Developmental plasticity of upper lethal temperature trait

To assess whether developmental temperature influences thermal tolerance, we conducted a split-brood experiment and performed a ramping assay to determine the upper lethal temperature of adult copepods reared at ambient (18°C) versus elevated (22°C) temperatures. Copepods that developed for one generation under 22°C exhibited an average ULT of 37°C and the average ULT of animals that developed at 18°C was 35.5°C. This 1.5°C difference in upper lethal temperature was found to be significant (*F* = 36.65; *P*-value= .00001) (Fig. 1B).

### Global differential gene expression analysis

We tested for differential gene expression between and within developmental temperature treatments at the three final temperature experiences. After cleaning and filtering for quality, there was an average of 18.5 million reads per sample. After filtering for depth of coverage per gene, we had high-quality expression data for 25,527 genes. Principal component analysis revealed strong effects of both developmental (PC1) explaining 38% of the variance and final temperature (PC2) explaining 19% of the variance on gene expression (Fig. 2). In addition, development at 22°C induced greater variability in gene expression, as reflected in the broader dispersion of samples in PCA space compared to those developed at 18°C (Fig. 2).

**Figure 2.**
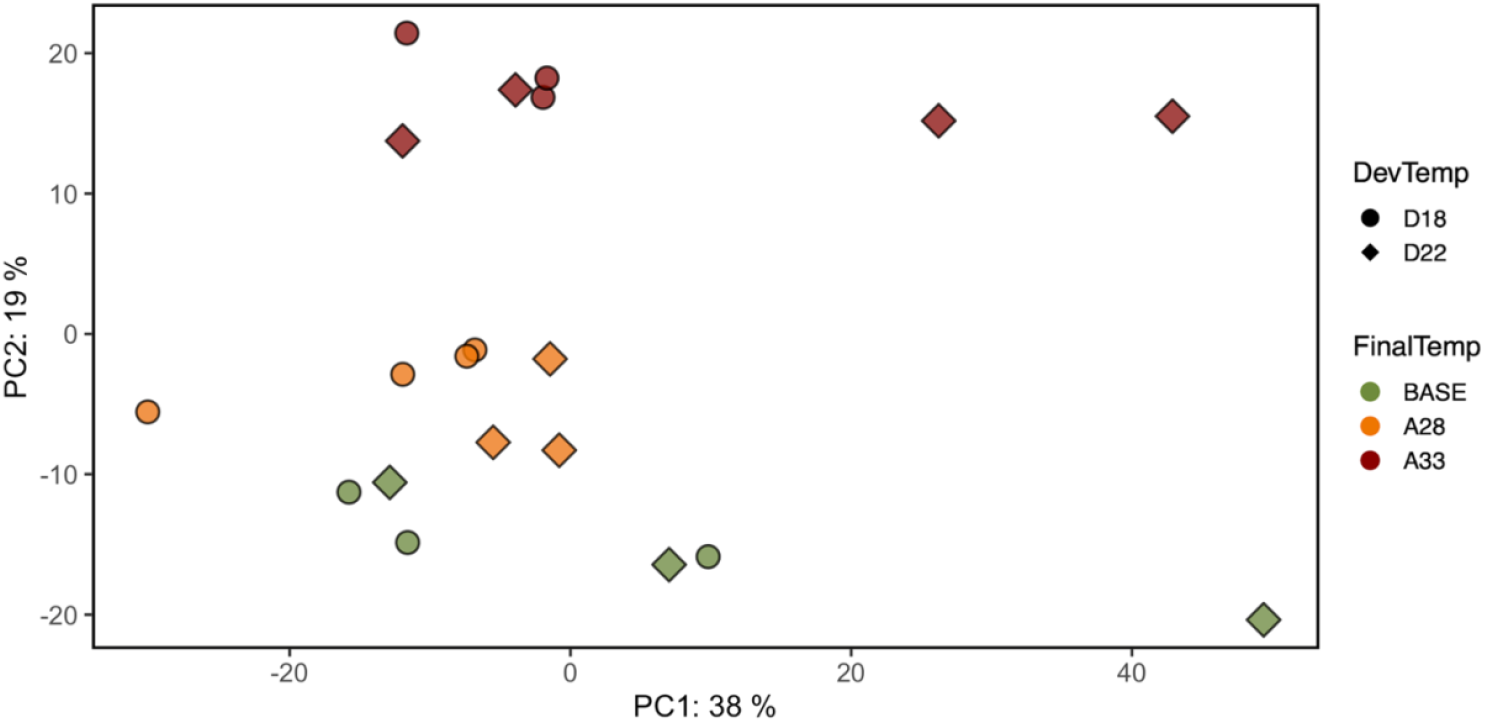
Principal component analysis of all genes across all samples. Developmental temperature (DevTemp) is indicated by point shape while Final temperature (FinalTemp) is indicated by point color and hue. Baseline (BASE) samples were collected at respective developmental temperatures 18°C and 22°C while acute exposures experienced one hour at 28°C (A28) or 33°C (A33).

### Differential gene expression between developmental temperatures

To examine how developmental temperature influenced gene expression responses to acute heat exposure, we compared transcriptomic profiles between copepods reared at 18°C versus 22°C across all final temperatures. The directionality and significance of differential gene expression between developmental temperatures at each final temperature were visualized using volcano plots (Fig. 3A-C). Each contrast showed higher relative expression in animals that developed at 22 compared to 18 for all three final temperatures (red points, Fig. 3A-C). The greatest upregulation in terms of magnitude of expression and number of genes occurred at 28°C with two and four times as many differentially expressed genes compared to differences at baseline and 33°C, respectively, with 233 DEGs at baseline, 482 at 28°C, and 104 at 33°C (Fig. 3A-C), indicating that the greatest differences in physiological responses between developmental temperatures was in response to this intermediate thermal stress, one hour at 28°C. Strikingly, the response to the highest temperature (33°C) had the most shared response between developmental temperatures (Fig. 3D).

**Figure 3.**
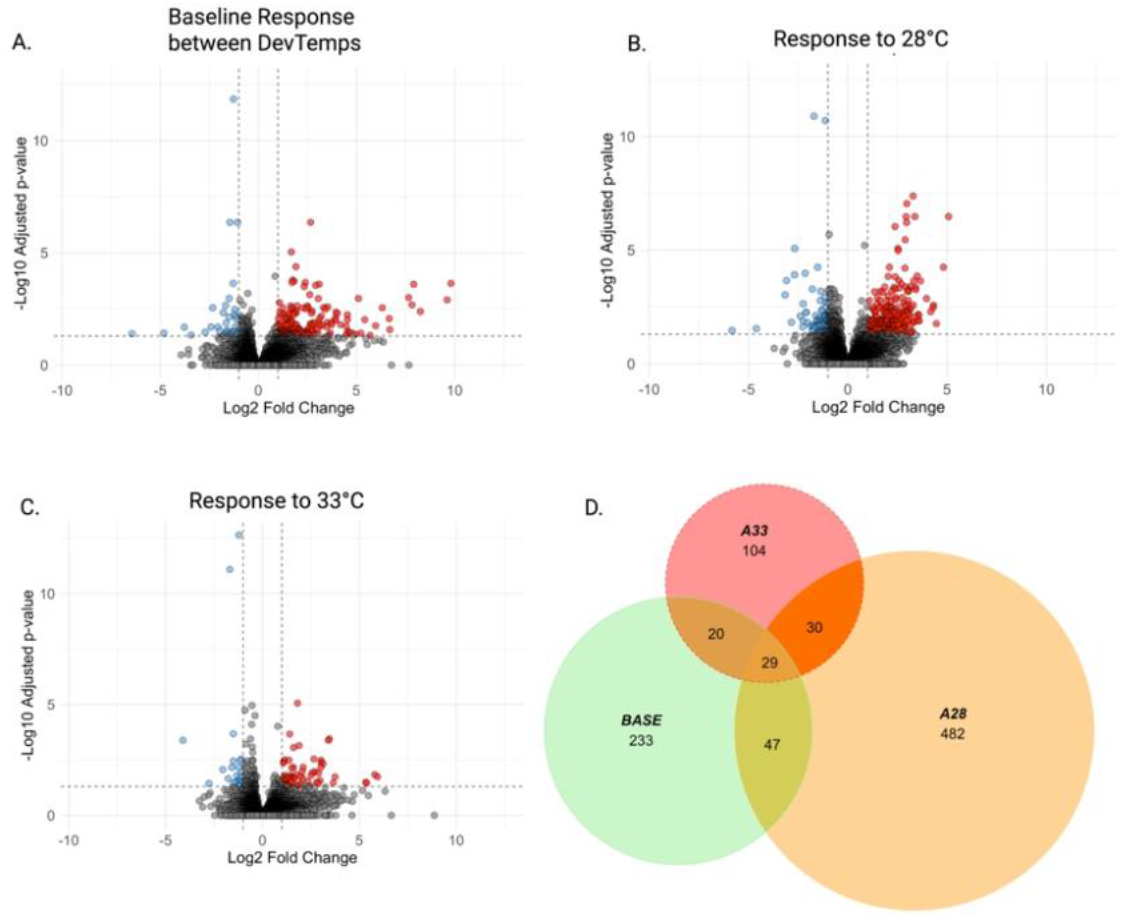
Differential gene expression between developmental temperatures at three final temperature conditions. (A–C) Volcano plots showing gene expression differences between copepods developed at 18°C (DT18) and 22°C (DT22) after exposure to baseline (A), 28°C (B), or 33°C (C). Each point represents a gene, with log_2_ fold change on the x-axis and –log_10_ adjusted p-value on the y-axis. Genes significantly upregulated in DT22 are shown in red, those upregulated in DT18 in blue, and non-significant genes in gray (adjusted p < 0.05). (D) Euler plot comparing the number of differentially expressed genes (DEGs) between developmental temperatures at each final temperature. Green represents DEGs at baseline, orange at 28°C, and red at 33°C.

To understand responses to acute temperature challenges within each developmental temperature, we compared gene expression within each developmental temperature between baseline and acute final temperatures. In contrast to the above results, within a developmental temperature, there was a limited transcriptional response between baseline and 28°C but a strong upregulation response between baseline and 33°C (Fig. 4). This was true for both development at 18°C (Fig. 4A-C) and development at 22°C (Fig. 4D-F). Euler plots highlight that the responses to these two sublethal temperature challenges were largely shared within each developmental temperature (Fig. 4C & 4E), with only 11 DEGs uniquely differentially expressed between baseline and 28°C at DT18 and 9 DEGs uniquely differentially expressed between baseline and 28°C at DT22. These results indicate that while exposure to 28°C induces minimal transcriptional changes, the transition to 33°C elicits a strong, and largely shared (Fig. 4C & F), gene expression response within both developmental temperatures.

**Figure 4.**
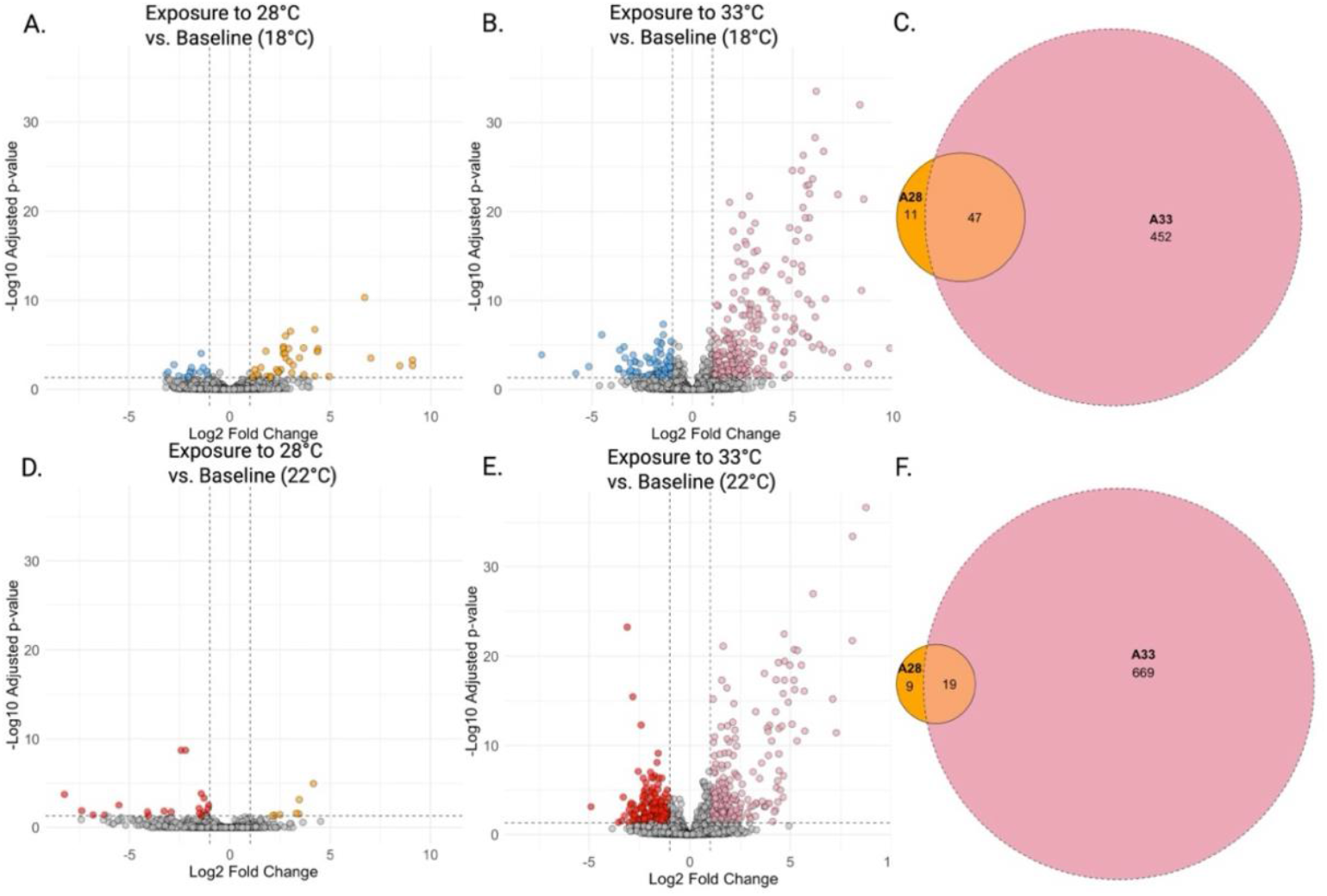
Differential gene expression between baseline and acute temperature exposures within each developmental temperature. Volcano plots show differentially expressed genes (DEGs) in copepods developed at 18°C (top row) and 22°C (bottom row) following 1-hour acute exposure and compared to baseline gene expression. Genes that were significantly upregulated in response to 28°C are shown in orange (A and D). Genes that were significantly upregulated in response to 33°C are shown in pink in (B and E). Each point represents a gene, with log_2_ fold change on the x-axis and –log_10_ adjusted p-value on the y-axis. Non-significant genes are shown gray (adjusted p < 0.05). Euler plots (C and F) summarize the number of DEGs within each developmental temperature group. Orange indicates DEGs between baseline and 28°C exposure, and pink indicates DEGs between baseline and 33°C exposure.

### Comparative gene expression at acute high temperature

To understand how developmental temperature affected acute thermal stress response, for each acute temperature, we plotted the log2-fold-change responses to that temperature versus baseline for animals developed at 18°C against the log2-fold-change responses to that temperature versus baseline for animals developed at 22°C (Fig. 5). Each point represents a gene, colored by significance category: genes significantly differentially expressed in both developmental treatments, in D18 only, or in D22 only (Fig. 5). In response to 28°C, 6 genes were differentially expressed in both contrasts, baseline vs. 28°C for each developmental temperature, 45 in D18 only, and 22 in D22 only (Fig. 5A). In contrast, in response to 33°C, 279 were differentially expressed in both contrasts, baseline vs. 33°C for each developmental temperature, 215 in D18 only, and 409 in D22 only. These plots illustrate a much stronger shared transcriptional response to the higher acute temperature of 33°C (Fig. 5B) than in response to 28°C across developmental temperatures (chi-square = 730.22, df = 3, *P* < 0.0001). Similarly, linear regression of log_2_ fold changes of all genes at DT18 vs. DT22 copepods for each acute exposure showed a much stronger correlation in response to 33°C (R^2^ = 0.252; p-value < 2.2e-16; Fig. 5B) than in response to 28°C (R^2^ = 0.047; *P* < 2.2e-16; Fig. 5A), suggesting that both developmental groups activated a more similar transcriptional response under extreme heat stress but with an overall stronger response from animals developed at 22°C (red points in Fig. 5B).

**Figure 5.**
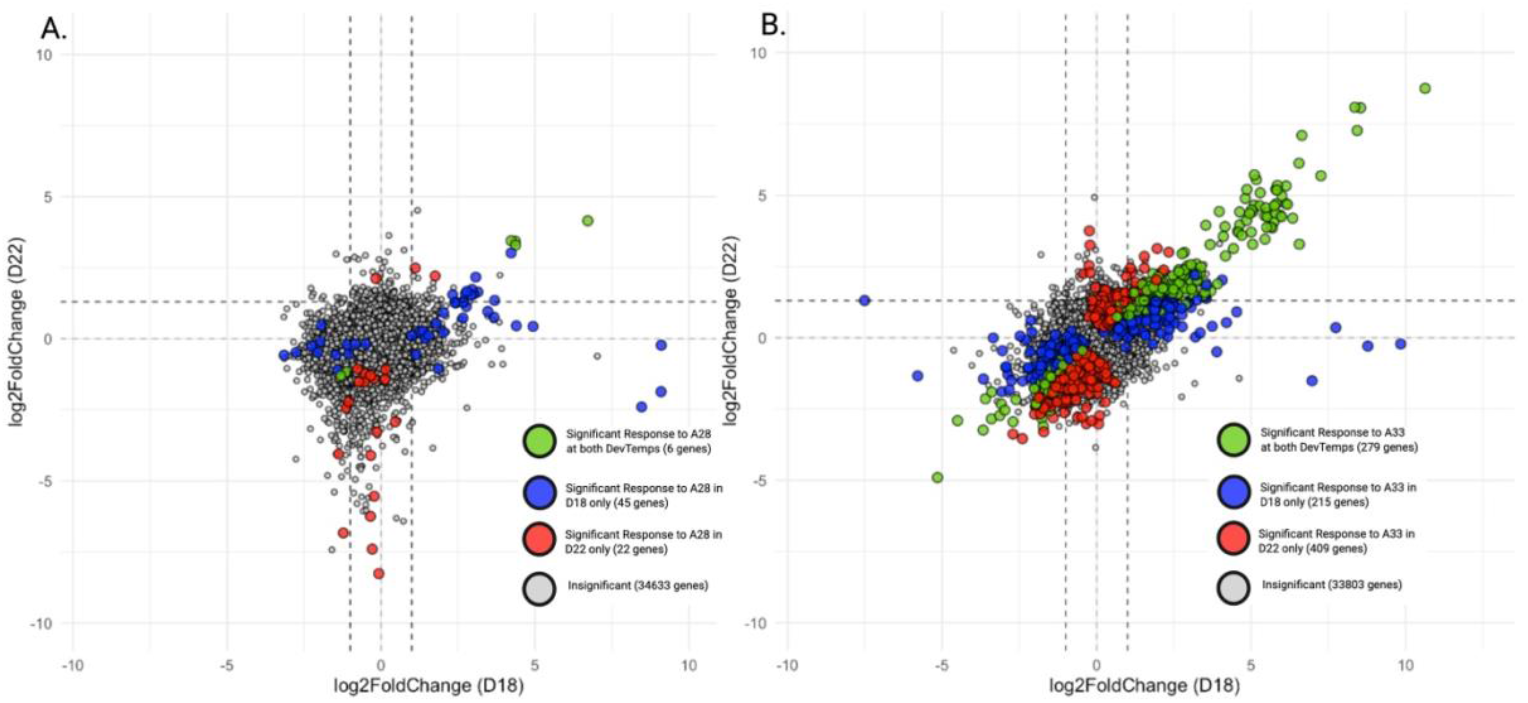
Comparison of gene expression responses to acute heat exposure at 28°C (A) and 33°C (B) for copepods reared at 18°C (D18) versus 22°C (D22). Each point represents a gene, with axes indicating the log2 fold change in expression in response to acute temperature exposure relative to baseline conditions for each developmental treatment. Genes are colored based on statistical significance: green = significant in both D18 and D22, blue = significant in D18 only, red = significant in D22 only, and gray = not significant in either developmental treatment. Dashed lines indicate zero log2 fold change.

### Gene ontology (GO) functional analysis

We performed Gene Ontology (GO) enrichment analysis to identify biological processes associated with differentially expressed genes (DEGs) across developmental and acute temperature conditions. We compared differential gene expression between developmental temperature treatments at each final temperature, as well as within developmental temperature treatments at baseline and each acute temperature treatment to gain a functional understanding of what types of genes were differentially responsive. GO categories enriched for the contrast between developmental temperatures at 28°C were particularly interesting given the strong divergent transcriptional response to that acute temperature. Our GO enrichment analysis revealed that key biological processes, including mRNA processing, negative regulation of metabolism, ribonucleoprotein-related, and cell growth regulation were among the most significantly enriched categories, particularly in comparing response to acute exposure to 28°C between developmental temperatures (Fig. 6).

**Figure 6.**
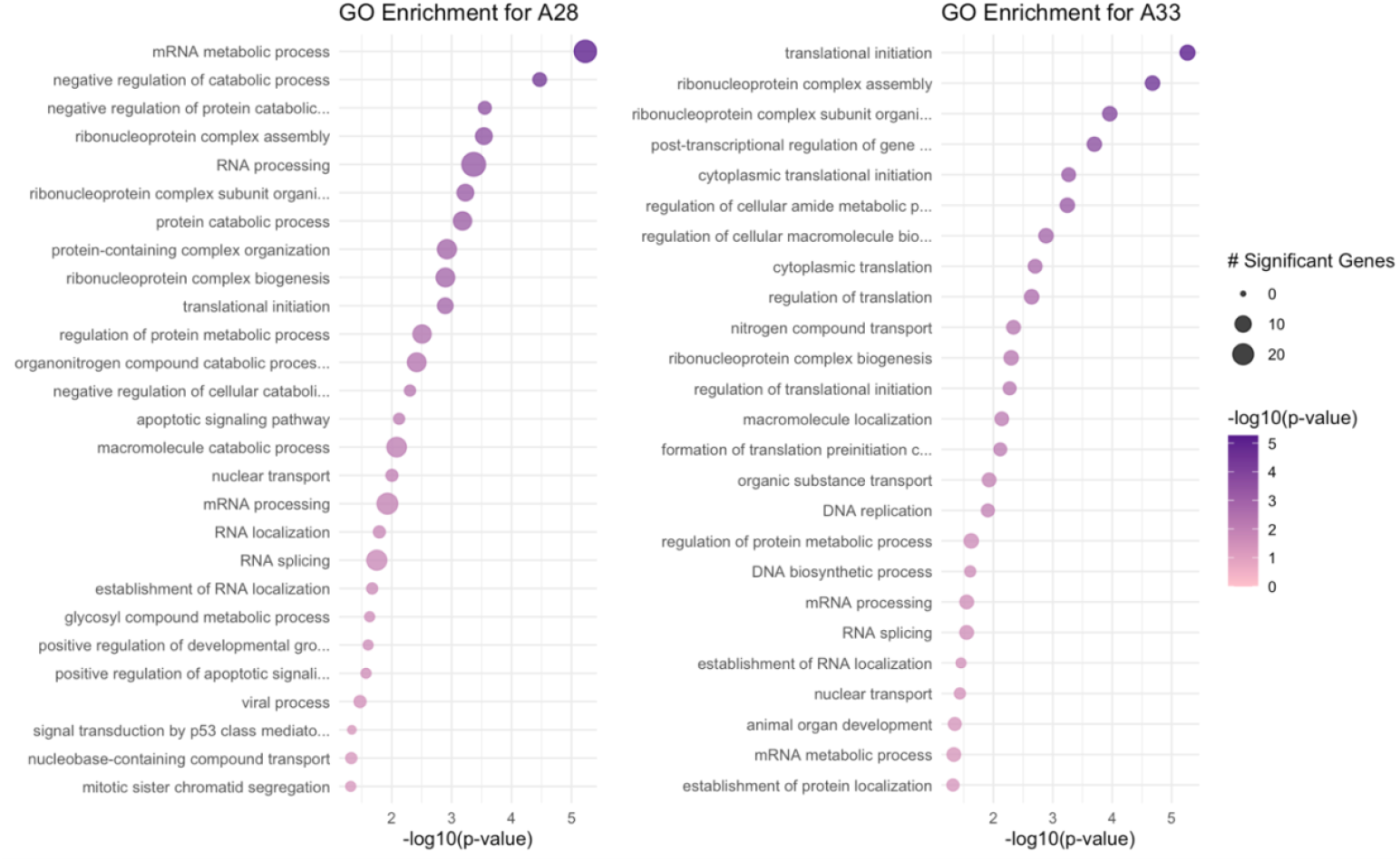
GO enrichment analysis of differentially expressed genes between developmental temperatures at 28°C and 33°C. The plot displays the enriched biological processes (BP) for genes significantly up- or downregulated between developmental temperatures at 28°C and 33°C. The x-axis represents the −log10(p-value) from the Fisher’s exact test, with larger values indicating stronger statistical significance. Dot size corresponds to the number of significant genes associated with each GO term, while color intensity (from pink to purple) reflects the −log10(p-value).

### Gene Co-Expression Networks to Identify Molecular Underpinnings of Thermal Tolerance

To investigate gene networks associated with thermal tolerance, we performed weighted gene co-expression network analysis (WGCNA) to identify modules of co-expressed genes and test for their association with the upper lethal temperature (ULT) phenotype. We identified 27 co-expression modules, three of which were correlated with ULT (*P* < 0.05; S1). The *green* module exhibited the strongest positive correlation with ULT (Pearson’s *r* = 0.89), with genes in this module showing increased expression in individuals with higher thermal tolerance. Conversely, the *darkgrey* module was strongly negatively correlated with ULT (*r* = −0.85), with genes in this module exhibiting lower expression in individuals with higher ULT. Boxplots illustrate differences in module expression across developmental and acute temperature treatments for both the positively correlated (Fig. 7A) and negatively correlated (Fig. 7B) modules, showing clear shifts in expression between copepods developed at 18°C and 22°C. Interestingly, for genes in the module positively correlated with ULT, the expression differences between developmental groups persisted at acute temperatures (Fig. 7A), but not for genes in the *negatively correlated module* module (Fig. 7B). In addition, scatterplots of module eigengene values plotted against ULT data from each sample further reveal the relationship between the network of a co-expressed genes that the thermal phenotype: the eigengene values of the positively correlated module increased with higher ULT, i.e., gene expression for genes in this module were positively correlated with ULT phenotypic variation across samples (Fig. 7C). In contrast, the eigengene values of the negatively correlated module decreased with higher ULT, i.e., gene expression for genes in this module were negatively correlated with ULT phenotypic variation across samples (Fig. 7D). Together, these results demonstrate that modules positively and negatively associated with thermal tolerance show distinct patterns of developmental regulation and trait correlation.

**Figure 7:**
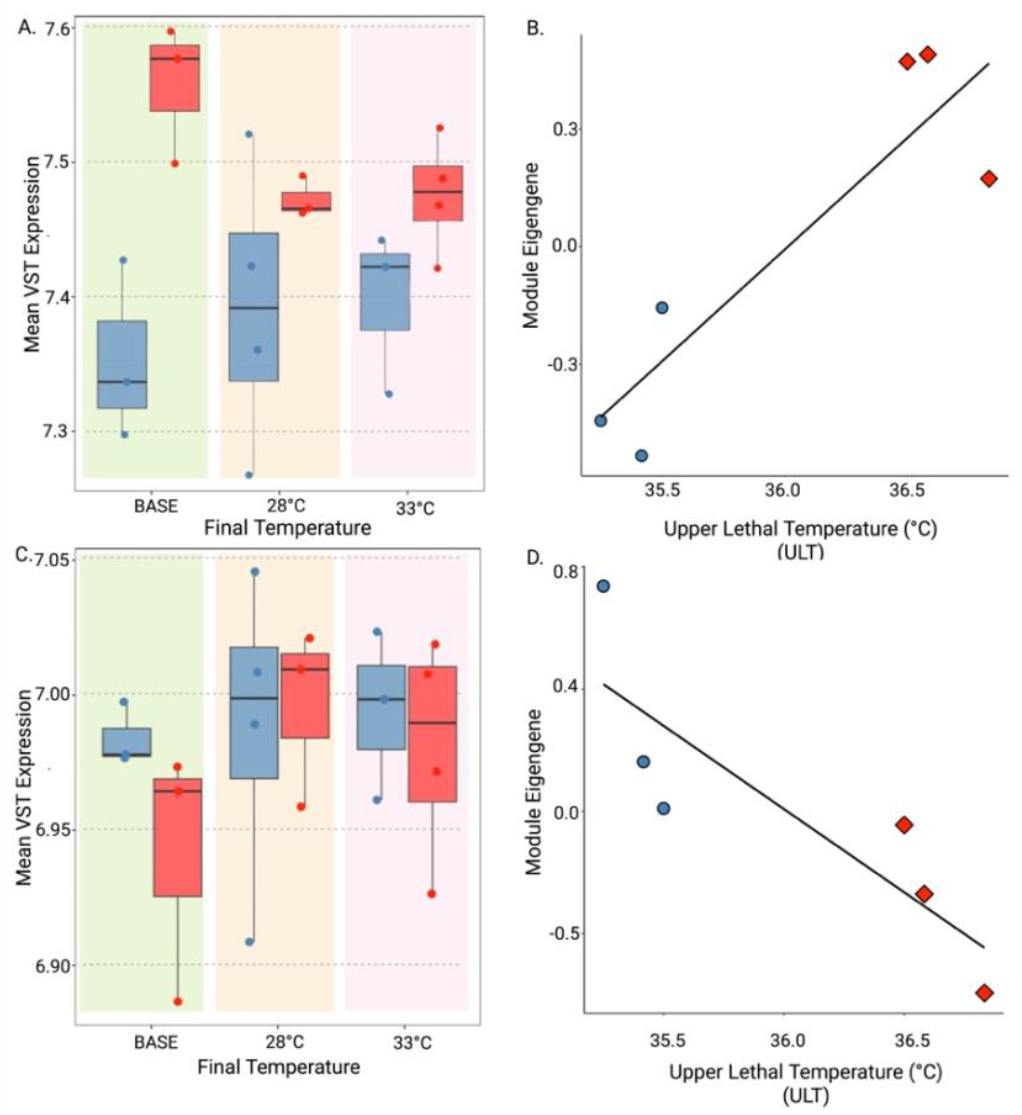
WGCNA module expression for modules significantly correlated with thermal tolerance trait. Boxplots show the mean variance-stabilized expression of all genes in the module most correlated with ULT (A) and negatively correlated with ULT (C) grouped by developmental temperature (18°C = blue, 22°C = red) and final acute temperature exposure (BASE (green), 28°C (orange), and 33°C (pink). Each point represents one sample. Scatterplots show module eigengene values (the first principal component summarizing module expression) plotted against upper lethal temperature (ULT) for each sample in the module most positively correlated with ULT (B) and negatively correlated with ULT (D). The shapes represent developmental treatment groups blue circles are samples that developed at 18°C and red diamonds are samples that developed at 22°C. Regression lines indicate module-specific associations with ULT.

To further understand the genomic bases underlying differences in thermal tolerance at baseline, we subset genes that were both differentially expressed at baseline between developmental temperatures and present in the module that is most highly positively correlated with increased upper lethal temperature. By focusing on genes that were both differentially expressed at baseline and associated with ULT, we aimed to identify key biological processes that may contribute to thermal tolerance. We found that the 24 genes in this group were involved in several key biological processes related to stress response, energy metabolism, gene regulation, and cellular homeostasis. Specifically in this group, we saw the genes adenosine kinase and cAMP dependent kinase which are involved in cellular signaling and metabolic regulation; RNA-binding lark, RNA polymerase I, and T-complex I which are involved in gene regulation and protein homeostasis; and oxidoreductase HTATIP2, amine oxidase, Vonwillebrand factor, and chitinase which are involved in stress response and cellular defense.

## Discussion

Our results demonstrate that developmental temperature leads to significant differences in thermal tolerance, with copepods reared at a warmer temperature exhibiting higher upper lethal temperatures than those developed at a lower temperature. This is consistent with previous findings of developmental plasticity in *Acartia tonsa* and suggests that early-life thermal environments can prime physiological responses to future stress. To investigate the molecular bases of this plasticity, we examined gene expression patterns shaped by both developmental history and the severity of acute temperature exposure. We found that moderate acute temperatures (28°C) revealed divergent transcriptional responses between developmental treatments, while higher acute temperatures (33°C) induced a more conserved expression profile across developmental groups. This pattern is consistent with the concept of transcriptional front-loading, where elevated baseline expression of stress-related genes in individuals reared at warmer developmental temperatures may contribute to enhanced resilience (Barshis et al., 2013). Gene co-expression network analysis further identified modules associated with thermal tolerance, highlighting key functional categories including stress response, energy metabolism, and gene regulation. Together, these findings support a model in which the developmental environment shapes transcriptional responses, contributing to plasticity in gene expression and thermal tolerance.

### Moderate thermal stress reveals developmentally programmed transcriptomic differences while higher heat induces a convergent, conserved cellular response

A key novel aspect of our study is the explicit comparison of transcriptomic responses to both moderate and extreme acute stress levels. Our design allowed us to test whether the severity of thermal stress influences the degree to which responses are developmentally programmed or converge on a conserved cellular strategy. By including both 28°C and 33°C challenges, we demonstrate that moderate stress amplifies developmental differences in gene regulation, whereas extreme stress overrides these differences, driving a shared stress response. This comparative framework provides new evidence that the nature of plasticity itself is conditional on the intensity of environmental stress.

Our results reveal that developmental temperature drives distinct transcriptomic responses to short-term increased temperature, which may underlie developmentally plasticity in thermal tolerance. When comparing gene expression between developmental temperatures at each final temperature and within each developmental temperature, we observed the greatest number of differentially expressed genes (DEGs) at 28°C between developmental groups (Fig. 2). GO analysis revealed that the genes differentially expressed between DT18 and DT22 at 28°C were enriched for mRNA metabolic processes and RNA processing (Fig. 6). This enrichment highlights how the regulation of these processes may act as central mechanisms underlying the plastic response to sub-lethal heat stress. The large number of differentially expressed genes suggests that a moderate temperature increase amplifies molecular differences between developmental treatment groups, potentially reflecting a front-loaded transcriptional response in which copepods reared at 22°C upregulate key stress and regulatory pathways in anticipation of future thermal stress (Fig. 3). Upregulation of these genes may reflect a preparatory response within these important pathways in copepods reared at 22°C that contributes to their higher thermal tolerance (Cavieres & Sabat, 2008). This phenomenon has been observed in corals from variable thermal environments, where constitutively higher expression of stress-related genes confers greater resilience to heat exposure, and may represent a common strategy among marine ectotherms facing environmental variability (Barshis et al., 2013). Similar patterns have been documented more broadly: for instance, warm acclimation in aquatic invertebrates led to elevated baseline expression of stress, immune and protein-synthesis genes that reduced the need for further induction upon exposure to hypoxia, facilitating cross-tolerance between stressors (Collins et al., 2021). In terrestrial systems, among multiple *Acacia* tree species adapted to different thermal regimes, early transcriptional heat responses were significantly associated with the species’ native climates, suggesting that frontloading of temperature-responsive genes may be a widespread adaptive mechanism (Andrew et al., 2024). These examples across marine and terrestrial taxa suggest that anticipatory upregulation of stress pathways may be a broadly conserved mechanism that enhances resilience.

In contrast, we observed fewer DEGs between developmental temperature groups at 33 °C, suggesting a convergent stress response in which both groups rely on a shared set of heat shock and stress-related genes (Fig. 3). This pattern mirrors findings in other systems where moderate stress reveals developmental or population-level differences, but extreme stress induces a more uniform cellular response. In *Drosophila melanogaster*, mild-to-moderate heat stress elicited distinct transcriptomic profiles between developmental treatments, whereas severe heat exposure produced conserved expression patterns across groups (Lukoszek et al., 2016). Similarly, in the marine copepod *Temora longicornis*, transcriptomic divergence between thermal histories was greatest under moderate warming but diminished under more extreme stress, as both groups converged on a core set of stress-related genes (Semmouri et al., 2019). These parallels suggest that while developmental programming can shape gene expression strategies under moderate stress, approaching physiological limits constrains transcriptomic flexibility, leading to reliance on conserved protective mechanisms.

Although both developmental treatment groups exhibited substantial gene expression responses to 28°C, the sets of responsive genes differed between the groups, indicating distinct genomic strategies for coping with thermal stress. When comparing the gene expression response for each developmental group between baseline (18°C or 22°C) vs moderate heat challenge (28°C), there were only six genes in common though dozens were uniquely regulated in each group (Fig. 6). In animals at DT18, we observed HSP70 and HSP105 to be among the top three differentially expressed genes at 28°C compared to baseline suggesting that copepods reared in cooler conditions mounted a more reactive heat shock response to 28°C. Interestingly, the same heat-shock genes were not significantly differentially regulated in DT22 animals between baseline and 28 °C—but they were among the top differentially expressed genes in DT22 animals exposed to 33 °C, suggesting that developmental at a warmer temperature primes the copepods to tolerate moderate heat without triggering the heat shock pathway, effectively raising the activation threshold. This pattern implies that warmer developmental temperatures may confer thermal preconditioning that delays heat shock gene activation until more stressful conditions occur (Schoville et al., 2012; Hoffman et al., 2024).

This work suggests that copepods reared in warmer conditions may require more extreme heat to activate the heat shock pathway, reflecting a primed but threshold-dependent response, while those developed in cooler conditions activate stress pathways more readily under moderate challenge, but the more constrained plasticity may lack sufficient capacity or flexibility under extreme heat. These findings support prior work that observed that *Acartia tonsa* from cooler environments activate stress responses at a lower heat threshold, whereas individuals acclimated to a warmer environment for 24-hours require higher temperatures to induce activation (Rahlff et al., 2017). Similarly, work in tropical estuarine copepods demonstrated that delayed induction of metabolic stress and HSP expression until they approached upper thermal limits (Low et al., 2018). These studies support the idea that thermal history sets response thresholds, aligning with our observed heat shock expression asymmetry between DT18 and DT22. This divergence is further supported by regression analysis of gene-level expression changes at 28°C, which revealed a weak but significant relationship between DT18 and DT22 fold changes (R^2^ = 0.047; p-value < 2.2e-16; Fig. 5A), indicating limited coordination in transcriptional responses between developmental treatments. At 33°C, however, transcriptional responses between developmental groups were more aligned. At 33 °C, approximately 30% of all significant differentially expressed genes (DEGs) were shared between developmental temperature treatments, whereas at 28°C only 8% were shared, indicating that transcriptional responses to acute stress become more convergent under extreme heat exposure. The scatter plot revealed a tighter clustering of genes along the diagonal, suggesting similar expression patterns between DT18 and DT 22 (R^2^=0.252; p-value < 2.2e-16; Fig. 5B).

Our work aligns with broader theories of stress-induced plasticity, which posit that plasticity is greatest under moderate conditions and becomes constrained as organisms approach physiological limits (Tomanek & Zuzow, 2010). At 28°C, copepods experience a moderate heat stress condition requiring a more flexible, plastic response, whereas 33°C may push copepods closer to their physiological limits, leading to a more constrained gene expression profile dominated by conserved stress responses. GO analysis of genes differentially expressed between baseline and 33°C within each developmental temperature reveals enrichment for canonical stress response categories, including cell death, ubiquitin regulation, and heat response. At the higher acute temperature, regardless of developmental temperature, the stress response becomes more uniform and limited to essential protective pathways, a pattern consistent with findings in *Drosophila melanogaster* embryos, where conserved stress responses dominate under extreme heat stress (Lockwood et al., 2010; Mikucki et al., 2024). This shared response is reflected in the smaller number of DEGs between developmental temperatures at 33°C compared to 28°C (Fig. 3D). Taken together, these results align with the idea that organisms can fine-tune their gene expression under moderate stress but rely on core protective mechanisms under extreme conditions (King & Stillman, 2022). This finding is consistent with the broader concept that phenotypic plasticity is highest under intermediate stress levels, whereas extreme stress constrains molecular responses to essential survival mechanisms (Logan & Buckley, 2015). Developmental temperature may further shape this response by shifting the thresholds at which plasticity is activated, with organisms reared at warmer temperatures exhibit greater thermal tolerance but potentially reduced plasticity at higher temperatures (van Heerwaarden et al., 2024).

### Developmental Programming Enhances Expression of Genes Linked to Thermal Tolerance

We found that genes whose expression was highly correlated with thermal tolerance were differentially regulated between baseline developmental temperature treatments, indicating that regulation of these genes may underlie developmental plasticity. Genes in the module most positively correlated with ULT, exhibited higher expression in copepods developed at 22°C, suggesting that in response to a warmer rearing temperature these genes were upregulated and that these genes were correlated with improved thermal tolerance (Fig. 7A). In contrast genes in the module most negatively correlated with ULT were expressed at higher levels in copepods developed at 18°C, indicating that these individuals rely more on pathways that were downregulated in copepods with higher thermal tolerance (Fig. 7C). This pattern of molecular preconditioning aligns with other studies of thermal plasticity, where organisms reared in warmer environments exhibit elevated expression of heat-responsive genes and enhanced heat tolerance (Ghalambor et al., 2015; Havird et al., 2020), suggesting that copepods reared at 22°C display a form of anticipatory plasticity. Such transcriptional shifts likely underlie phenotypic plasticity of thermal tolerance provide a rapid and flexible response to environmental change, especially in fluctuating environments where stress timing and magnitude are unpredictable.

Genes that were both differentially expressed at baseline between developmental temperatures and positively correlated with upper lethal temperature likely directly contribute to thermal tolerance. We identified 24 such genes which spanned several essential biological functions, including stress response, energy metabolism, gene regulation, and cellular homeostasis. These functional categories are consistent with key physiological processes known to underlie thermal resilience across ectothermic species (Somero, 2010). Notably, genes such as adenosine kinase and cAMP-dependent kinase play key roles in cellular signaling and energy metabolism, both of which are critical for maintaining physiological function under thermal stress (Feder & Hofmann, 1999; Schulte, 2015). Genes involved in gene regulation and protein stability—such as RNA-binding lark, RNA polymerase I, and T-complex protein I— highlight the importance of transcriptional and translational control in developmentally plastic responses to temperature (Tomanek, 2010; Evans & Hofmann, 2012). Additionally, stress-responsive genes such as HTATIP2 (a redox-sensitive oxidoreductase), amine oxidase, chitinase, and Von Willebrand factor-like protein likely contribute to cellular protection, damage repair, and redox balance under elevated temperature conditions (Kültz, 2005; Somero, 2010). These genes illustrate an integrated molecular toolkit that enables copepods reared at warmer temperatures to anticipate and mitigate the cellular challenges of elevated temperature.

Together, these results suggest that copepods reared at warmer developmental temperatures may exhibit a coordinated upregulation of both broad regulatory networks and specific functional genes contributing to thermal tolerance. The WGCNA results, which linked modules correlated with thermal tolerance to distinct gene expression patterns between developmental treatments, are consistent with transcriptional front-loading observed in other taxa. In reef-building corals, modules enriched for stress response genes with more highly expressed in heat tolerant individuals (Strader & Quigley, 2022) and similar patterns have been observed in mussels (Tomanek & Zuzow, 2010) and fish (Narum & Campbell, 2015). These parallels suggest that anticipatory regulation may be a conserved feature of plastic thermal response to environmental change.

### Developmental Plasticity May Support Resilience to Environmental Change

Our findings highlight the role of developmental plasticity in shaping copepod responses to heat stress. The ability of copepods to modulate their gene expression in response to developmental temperature may provide a mechanism for short-term acclimation to changing thermal regimes (Stillman, 2003). This type of phenotypic plasticity can enhance individual survival during environmental fluctuations by allowing organisms to adjust their molecular and physiological processes without requiring genetic changes. However, extreme environmental conditions may limit this plasticity, potentially constraining adaptive capacity under future climate scenarios (Huey et al., 2012). As temperatures continue to rise, organisms that rely heavily on phenotypic plasticity might face difficulties, as extreme stressors could overwhelm their capacity for flexible responses. The limitation of plasticity under extreme stress has been observed in various taxa, where organisms exhibiting greater flexibility at moderate temperatures show reduced plasticity under more severe conditions (Sih et al., 2011). This aligns with the conceptual framework that distinguishes adaptive from non-adaptive plasticity and highlights how plasticity may be insufficient to buffer organisms from rapid environmental change (Ghalambor et al., 2007).

Studies in other marine organisms suggest that populations with greater developmental plasticity may exhibit enhanced resilience to climate change, particularly in fluctuating environments (Kelly, 2019). In copepods, this plasticity could influence population persistence and distribution, enabling them to adapt to warming ocean temperatures and potentially novel temperature regimes. However, the success of such populations likely depends on the balance between plastic responses and genetic adaptation. The emergence of genetic adaptation, though slower than phenotypic plasticity, can provide long-term resilience (Ghalambor et al., 2007; Oostra et al., 2018; Stemkovski et al., 2025). Recent work in *Acartia tonsa* shows that populations undergoing long-term experimental evolution to warming conditions exhibited both initial plastic responses and later genetic adaptation, including a loss of transcriptional plasticity after adaptation had occurred. This finding suggests that plasticity may serve as a short-term buffer, while genetic change stabilizes traits under persistent environmental stress (Brennan, deMayo, Dam, Finiguerra, Baumann, & Pespeni, 2022). Whether plastic responses can persist across generations, and how they interact with or are replaced by adaptive evolution, remains a critical question particularly in the context of rapid environmental change (Shama et al., 2014; Donelan et al., 2020; Puy et al., 2022; Brennan et al., 2025).

### Limits of Plasticity: Future Warming May Outpace Coping Capacity Without Genetic Change

While our study provides insights into transcriptomic responses to thermal stress, additional research is needed to determine the long-term effects of developmental temperature on fitness and reproductive success. Our results enhance the understanding of the mechanisms behind developmental plasticity which enables copepods to cope with environmental change. The power of developmental plasticity of thermal in *Acartia tonsa* has been established in work that found that a single generation of development at a warmer temperature conferred the same increase in acute thermal tolerance as over 40 generations of selection under warming conditions (Ashlock et al., 2024). However, whether this plasticity is sufficient to buffer against the pace of climate change remains unclear (Stillman, 2003; Kelly, 2019). While phenotypic plasticity can enhance short-term survival, long-term persistence may depend on genetic adaptation or transgenerational plasticity (Vinton et al., 2022). Experimental evolution studies in *A. tonsa* have shown genetic adaption after 25 generations adaptation (Brennan, deMayo, Dam, Finiguerra, Baumann, Buffalo, et al., 2022), loss and recovery of fitness within three generations (Dam et al., 2021), and a loss of transcriptional plasticity after long-term adaptation (Brennan, deMayo, Dam, Finiguerra, Baumann, & Pespeni, 2022). In addition, recent work has revealed consistent epigenetic divergence across generations during experimental evolution which was positively associated with gene expression divergence and negatively related to genetic divergence (Brennan et al., 2025). These findings suggest independent and complementary roles for genetic and epigenetic variation in global change adaptation. However, the mechanisms driving rapid changes in the early generations that may influence thermal tolerance in future generations are unknown. Investigating whether these transcriptomic changes persist across generations or are heritable could provide key insights into the evolutionary potential of thermal tolerance in marine ectotherms.

## Conclusion

This study provides compelling evidence that developmental temperature influences both baseline gene expression and the plasticity of transcriptional responses to acute thermal exposure. By linking these molecular responses to thermal tolerance traits, we shed light on the genomic underpinnings of developmental plasticity in *Acartia tonsa*. We found that moderate warming revealed developmentally programmed transcriptomic differences, while more extreme warming elicited a conserved response—demonstrating that the nature of plasticity itself is conditional on the environment. We also identified specific gene modules whose baseline expression levels predict thermal tolerance, highlighting molecular pathways involved in stress response and gene regulation that may serve as targets of plasticity or selection under warming scenarios. These findings align with broader patterns observed in marine ectotherms, where transcriptional plasticity plays a pivotal role in coping with rapid environmental changes (Sunday et al., 2011; Rivera et al., 2021) and expands on this work by demonstrating that the magnitude and direction of transcriptomic responses vary depending on developmental temperature and intensity of short-term exposure conditions.

As global ocean temperatures continue to rise, understanding the mechanisms of thermal tolerance and plasticity will be essential for predicting species resilience. Further work in this area should explore the genetic and epigenetic mechanisms that modulate these observed differences in gene expression to determine whether these responses are heritable and how they might shape population persistence across generations. These findings suggest that while developmental plasticity supports short-term resilience, it may be insufficient under extreme stress. The interplay between developmental plasticity, genetic adaptation, and transgenerational effects will be central to understanding the long-term survival and distribution of species facing shifting climate patterns. By demonstrating how developmental temperature shapes both gene expression and phenotype, and identifying specific genes that contribute to thermal plasticity, our results offer novel insights into the mechanisms of short-term resilience and the potential targets of selection under future warming scenarios. The insights gained from this study contribute to a growing body of evidence supporting the role of developmental plasticity in mitigating the impacts of climate change on marine biodiversity.

## Acknowledgments

The authors acknowledge funding from the National Science Foundation Career Award 1943316 to MP. Discussions with Dr. M. Sasaki contributed to protocol design. The authors acknowledge the Vermont Advanced Computing Center (VACC) at the University of Vermont for providing computational resources that have contributed to the research results reported within this paper. URL: http://www.uvm.edu/vacc. Thank you to Dr. H. Dam and A. Salvemini for their knowledge of algal cultivation and for providing us with stock algae cultures to grow in our lab to feed to the copepods. We are grateful to Pespeni lab member A. McCraken for sharing R code for various steps in the bioinformatic analysis process.

## Data Accessibility

The data and code from this project are publicly available. The code is available on GitHub (https://github.com/aehall26/Atonsa_ThermalAcclimation). The gene counts, meta-data, and trait data are available on Dryad (DOI: 10.5061/dryad.866t1g23z). RNA sequencing data for each sample are available on the NCBI Sequence Read Archive (Accession number PRJNA1322529).

## Benefit-Sharing Statement

Benefits Generated: This research used copepod samples (Acartia tonsa) collected in U.S. coastal waters, where the Nagoya Protocol does not apply. Benefits from this research accrue from making all data and results publicly available through international repositories (NCBI SRA and Dryad, see Data Accessibility). More broadly, our findings contribute to the understanding of thermal tolerance and plasticity in marine organisms, with relevance to global climate change resilience and conservation.

## Author Contribution

All authors contributed to and approved the manuscript. AH and MP conceived and designed the study. AH and MRC performed the research, including culture maintenance, acute exposure experiments, and animal collection. AH performed RNA extractions. AH and MP performed data analysis. AH and MP created visualizations. AH and MP wrote and edited the manuscript.

