## Supplemental Figure 1 for "Developmental temperature drives distinct transcriptomic responses to acute temperatures and correlated differences in thermal tolerance"

MOLECULAR ECOLOGY

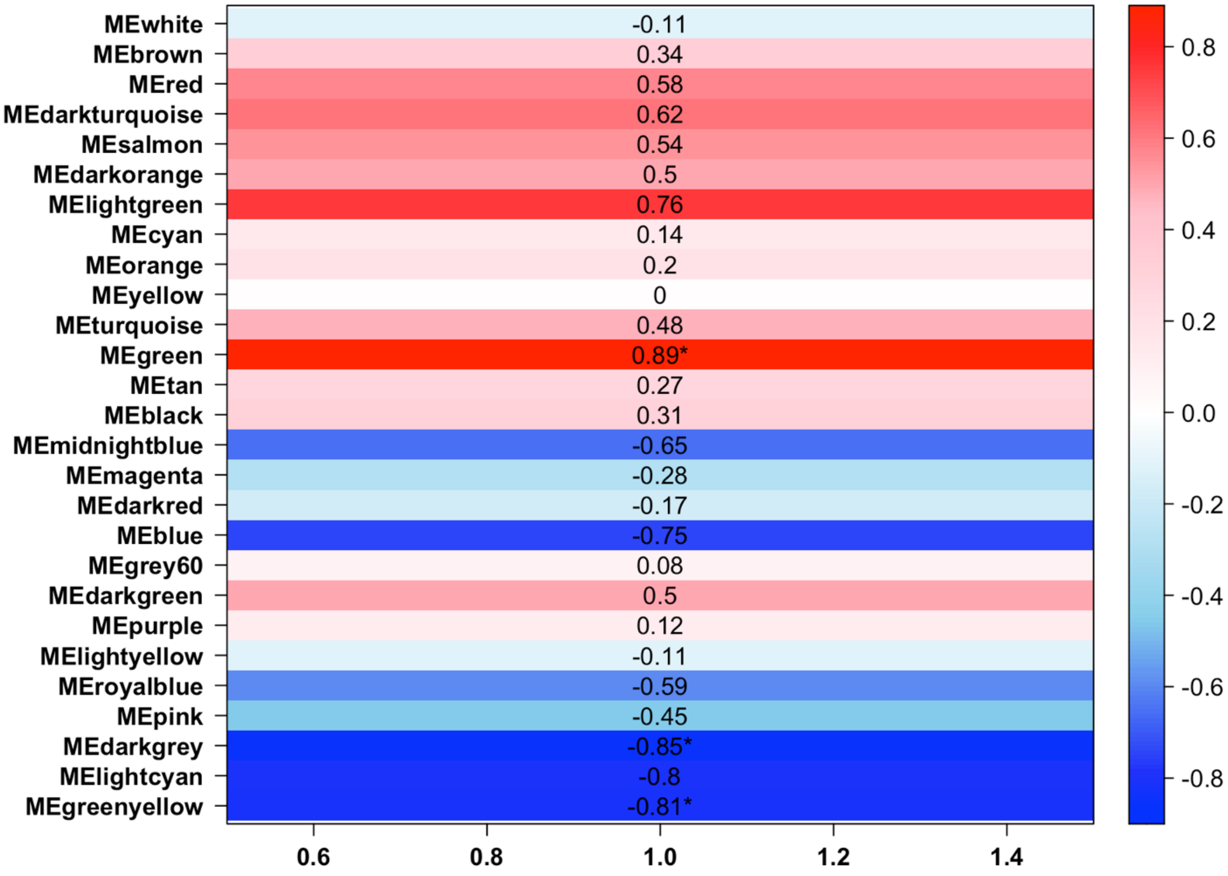


Supplemental Figure 1: **WGCNA module-trait correlations and gene expression heatmaps for the most correlated modules.** Module-trait correlations between WGCNA modules and upper lethal temperature (ULT). Each row represents a module eigengene (ME), with the corresponding Pearson correlation coefficient to ULT displayed. Positive correlations (red) indicate modules where gene expression increases with ULT, while negative correlations (blue) indicate modules where gene expression decreases with ULT. Asterisks denote statistically significant correlations.

MOLECULAR ECOLOGY
